# In-vivo estimation of axonal morphology from MRI and EEG data

**DOI:** 10.1101/2022.02.11.479116

**Authors:** Rita Oliveira, Andria Pelentritou, Giulia Di Domenicantonio, Marzia De Lucia, Antoine Lutti

## Abstract

**Purpose:** We present a novel approach that allows the estimation of morphological features of axonal fibers from data acquired in-vivo in humans. This approach allows the assessment of white matter microscopic properties non-invasively with improved specificity.

**Theory:** The proposed approach is based on a biophysical model of Magnetic Resonance Imaging (MRI) data and of axonal conduction velocity estimates obtained with Electroencephalography (EEG). In a white matter tract of interest, these data depend on 1) the distribution of axonal radius – *P*(*r*)– and 2) the g-ratio of the individual axons that compose this tract – *g*(*r*). *P*(*r*)is assumed to follow a Gamma distribution with mode and scale parameters, *M* and *θ*, and *g*(*r*) is described by a power-law with parameters *α* and *β*.

**Methods:** MRI and EEG data were recorded from 14 healthy volunteers. MRI data were collected with a 3T scanner. MRI g-ratio maps were computed and sampled along the visual transcallosal tract. EEG data were recorded using a 128-lead system with a visual Poffenberg paradigm. The interhemispheric transfer time and axonal conduction velocity were computed from the EEG current density at the group level. Using the MRI and EEG measures and the proposed model, we estimated morphological properties of axons in the visual transcallosal tract.

**Results:** The estimated interhemispheric transfer time was 11.72±2.87 ms, leading to an average conduction velocity across subjects of 13.22±1.18 m/s. Out of the 4 free parameters of the proposed model, we estimated *θ* – the width of the right tail of the axonal radius distribution and *β* – the scaling factor of the axonal g-ratio, a measure of fiber myelination. Across subjects, the parameter *θ* was 0.40±0.07 *µ*m and the parameter *β* was 0.67±0.02 *µm*^−*α*^.

**Conclusions:** The estimates of axonal radius and myelination are consistent with histological findings, illustrating the feasibility of this approach. The proposed method allows the measurement of the distribution of axonal radius and myelination within a white matter tract, opening new avenues for the combined study of brain structure and function, and for in-vivo histological studies of the human brain.

## 1 Introduction

The characterization of microscopic brain changes in-vivo in clinical populations is essential to the understanding of brain disease. Magnetic Resonance Imaging (MRI) is non-invasive and the primary technique for the assessment of brain structure in-vivo (Weiskopf et al., 2015; Kiselev and Novikov, 2018). MRI is sensitive to a large array of microscopic properties of brain tissue such as cell density, fiber radius and directionality, myelin, and iron concentration (Fukunaga et al., 2010; Lutti et al., 2014; Weiskopf et al., 2015; Jelescu et al., 2020). Biophysical models of the relationship between tissue microstructure and the MRI signal allow the assessment of microscopic properties of brain tissue from in-vivo MRI data (‘in-vivo histology’) (Weiskopf et al., 2015; Edwards et al., 2018; Kiselev and Novikov, 2018; Jelescu et al., 2020). Since MRI gives rise to a variety of image contrasts, each contrast being differentially sensitive to multiple microscopic properties of brain tissue, such biophysical models are intrinsically tuned to specific types of MR images (MacKay and Laule, 2016; Does, 2018; Kiselev and Novikov, 2018; Jelescu et al., 2020). Here, we focus on axonal radius and myelination and on the main biophysical models that allow their measurement from in-vivo data.

Axonal radius is a key property of neurons and is the main determinant of the speed of conduction of action potentials along axonal fibers (Rushton, 1951; Waxman and Bennett, 1972). Axonal radius plays an essential role in neuronal communications and is an instrumental structural underpinning of brain function (Liewald et al., 2014). Axonal radius estimates have been used as measures of functional connectivity in generative models of brain function, *e*.*g*. using dynamic causal modeling (Stephan et al., 2009; Honey et al., 2010). Moreover, axonal radius is a biomarker of brain development and healthy aging (Weiskopf et al., 2015) and is of high clinical relevance for a range of disorders such as autism (Wegiel et al., 2018), multiple sclerosis (Evangelou et al., 2001) and motor-neuron disease (Cluskey and Ramsden, 2001). Diffusion contrast is the main type of MRI data used for the measurement of axonal radius in-vivo. Suitable biophysical models include AxCaliber (Assaf et al., 2008; Barazany et al., 2009), which enables the estimation of the full distribution of axonal radius. ActiveAx is an alternative model that enables the estimation of axonal radius in all white matter tracts without *a priori* knowledge of fiber orientation (Alexander et al., 2010). However, this model provides a single summary index of axonal radius distribution, weighted towards larger axons (Jones et al., 2018; Veraart et al., 2020). Axonal radius estimates obtained in-vivo from diffusion MRI data are often overestimated compared to histological values (Aboitiz et al., 1992; Liewald et al., 2014) due to the limited gradient strength of MRI scanners and other confounding factors, such as the dominance of the extra-axonal signal (Burcaw et al., 2015; Nilsson et al., 2017; Jones et al., 2018; Lee et al., 2018; Jelescu et al., 2020; Veraart et al., 2020).

Besides axonal radius, axonal fiber myelination is also a crucial factor in the transmission of neuronal information and brain function (MacKay and Laule, 2016). The non-invasive assessment of myelination enables the study of brain plasticity in healthy individuals and brain changes in a range of neurological disorders (Lazari and Lipp, 2021). Relaxometry MRI data are in-vivo biomarkers of bulk myelin concentration within brain tissue (Lutti et al., 2014; Stüber et al., 2014). While their validity is supported by a large array of empirical evidence, these biomarkers lack an explicit link with the underlying histological properties of brain tissue, consequently hindering the interpretability of results (Weiskopf et al., 2015). Further specificity may be gained from MRI measures of the fraction of water embedded within the myelin sheath (Feintuch et al., 2007; MacKay and Laule, 2016; Does, 2018). Nonetheless, important aspects pertaining to exchange between compartments and suitable MRI acquisition sequences remain unclear (Dortch et al., 2013). Another effort towards improved specificity lies in MRI measures of the g-ratio, the relative thickness of the myelin sheath around axons (Stikov et al., 2011, 2015). MRI g-ratio estimates are aggregate measures of the axonal g-ratio, across all fibers present in each voxel of an MR image (West et al., 2016).

In summary, current MRI markers of axonal radius and fiber myelination are averages across populations of axons present in each voxel of an MR image. To date, estimating the distribution of these morphological features across axonal populations remain largely out of reach. To address this limitation, we propose a novel approach that enables the estimation, from in-vivo data, of the radius and myelination of axonal fibers, across the distribution of axonal populations in a white matter tract. This approach is based on the combination of electroencephalography (EEG) measures of signal conduction velocity along a white matter tract of interest, and of MRI measures of the g-ratio, sampled along the same tract. MRI and EEG have been jointly used in brain connectivity studies and for the combined study of brain structure and function (Westerhausen et al., 2006; Sui et al., 2014; Helbling et al., 2015; Horowitz et al., 2015; Deslauriers-Gauthier et al., 2019). In particular, the high temporal resolution of EEG allows the estimation of the interhemispheric transfer time (IHTT) (Saron and Davidson, 1989; Marzi, 1999) using the established visual Poffenberger paradigm (Westerhausen et al., 2006; Whitford et al., 2011; Friedrich et al., 2017; Chaumillon et al., 2018). Subsequently, an estimate of axonal conduction velocity can be computed (Caminiti et al., 2013; Horowitz et al., 2015). The dominant contributions of axonal radius and myelination to conduction velocity (Drakesmith et al., 2019) underline the complementarity of this EEG measure with the MRI measures described above. In the first part of this paper, we present the biophysical model underlying the proposed approach (Fig. 1). Numerical simulations are then conducted to illustrate the plausibility of our results in light of the histology literature and to assess the variability and accuracy of the morphological estimates. To illustrate the feasibility of the proposed approach, we present estimates of axonal morphology obtained using in-vivo data from the visual transcallosal white matter tract of healthy volunteers.

**Figure 1.**
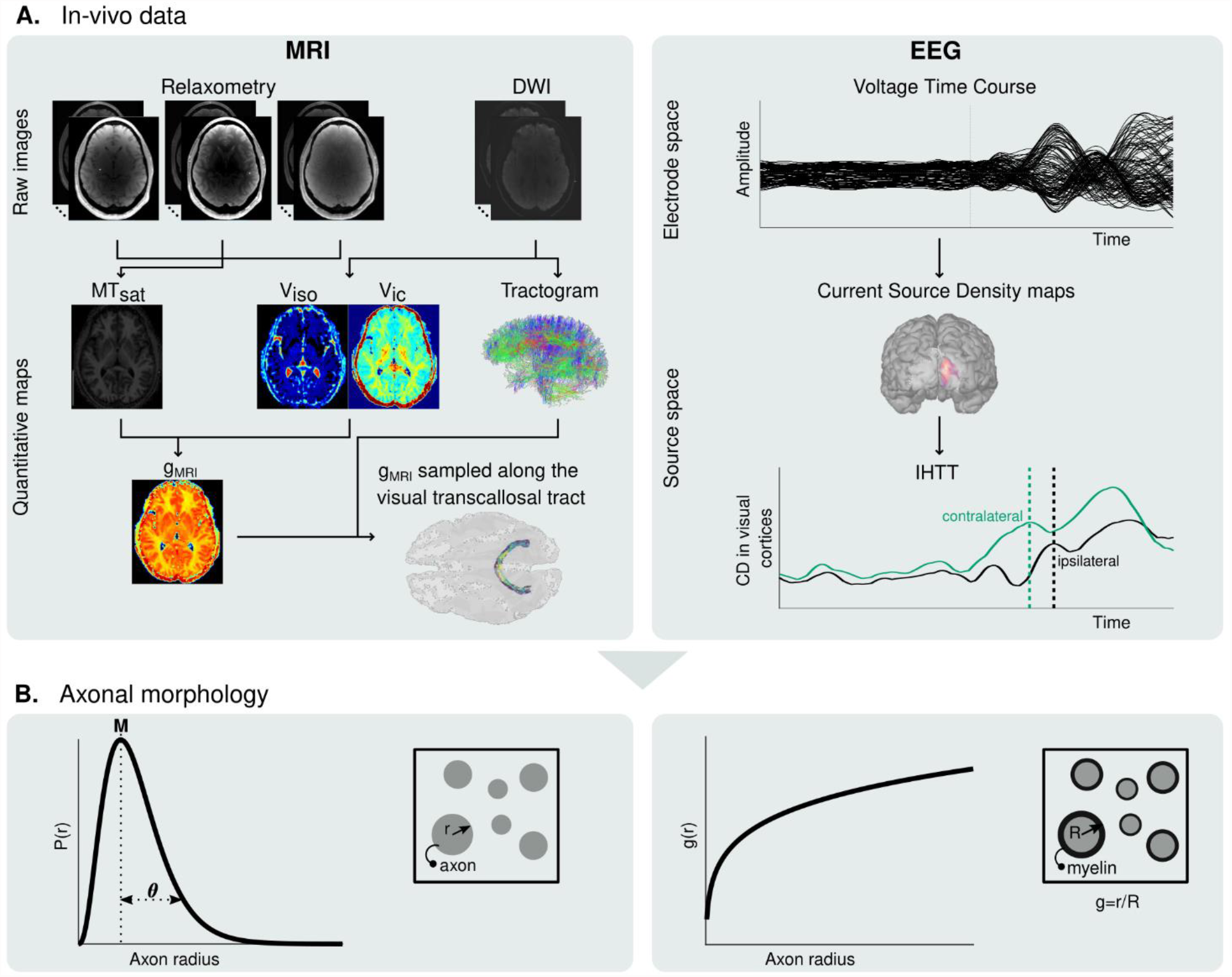
Estimation of microscopic morphological properties of axons from in-vivo MRI and EEG data. (A) Maps of MT_sat_, *v*_*iso*_ and *v*_*ic*_ are computed from the raw MRI data and subsequently used to compute maps of the MRI g-ratio (*g*_*MRI*_) (left). The MRI g-ratio maps are sampled along the visual transcallosal tract using the streamlines obtained with diffusion MRI tractography. Estimation of the IHTT is performed after source reconstruction of the EEG data, based on the difference in latency between the two maxima of activation in the two hemispheres observed on the group-averaged current source time course (right). (B) The *g*_*MRI*_ and IHTT estimates are used to estimate morphological properties of axons within the visual transcallosal tract (axonal radius distribution (*P*(*r*), left) and axonal g-ratio (*g*(*r*), right).

## 2 Theory - Biophysical model

### 2.1 Axon morphological properties

With the proposed model, both the MRI and EEG data are described as a function of the distribution of axonal radius (*P*(*r*)) and of the axonal g-ratio (*g*(*r*)) within a given white matter tract. The axonal radius distribution is assumed to be a Gamma distribution (Sepehrband et al., 2016):

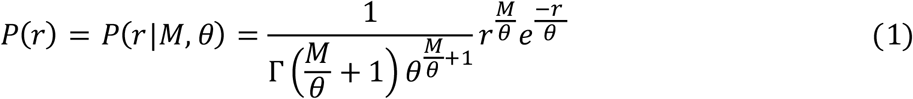

where *r* is the axonal radius. *M* represents the mode, *i*.*e*. the peak of the axonal radius distribution and the parameter *θ* represents the width of the right tail of the axonal radius distribution, a measure of the number of large axons in a white matter fiber tract.

Histological studies have shown that larger axons exhibit comparatively thinner myelin sheaths, *i*.*e*. larger g-ratios (Ikeda and Oka, 2012; Gibson et al., 2014). From data obtained in the peripheral nervous system of the rat, the radius dependence of the g-ratio was shown to follow (Ikeda and Oka, 2012): *g*_*REF*_(*r*) = 0.22 *log*(2*r*) + 0.508. However, to facilitate the mathematical manipulation of the biophysical model (Eqs. (3) and (5) below), we write the radius dependence of the axonal g-ratio as:

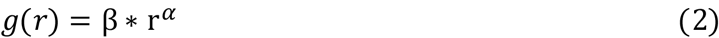

With this power-law, the exponent *α* represents the slope of the radius dependence of the axonal g-ratio, while the parameter *β* is a scaling factor. We found an excellent level of agreement between the power-law (Eq. (2)) and the reference relationship (*g*_*REF*_) of Ikeda and Oka, 2012 (R^2^=0.99), with *α*=0.18 and *β*=0.57 (Fig. 2A). Accounting for the higher g-ratio of axonal fibers in the human central nervous system (Aboitiz and Montiel, 2003; Caminiti et al., 2009; Liewald et al., 2014) requires the addition of a systematic offset to *g*_*REF*_ which we estimate at 0.14, assuming a g-ratio of 0.7 for an axonal radius of 0.9 *µ*m (Mohammadi et al., 2015; Stikov et al., 2015). However, the proposed power-law (Eq. (2)) does not explicitly include an offset term and its validity in the central nervous system should be verified. We found an excellent agreement with the reference relationship (*g*_*REF*_) of Ikeda and Oka (R^2^=0.99), with *α*=0.14 and *β*=0.71 (Fig. 2B).

**Figure 2.**
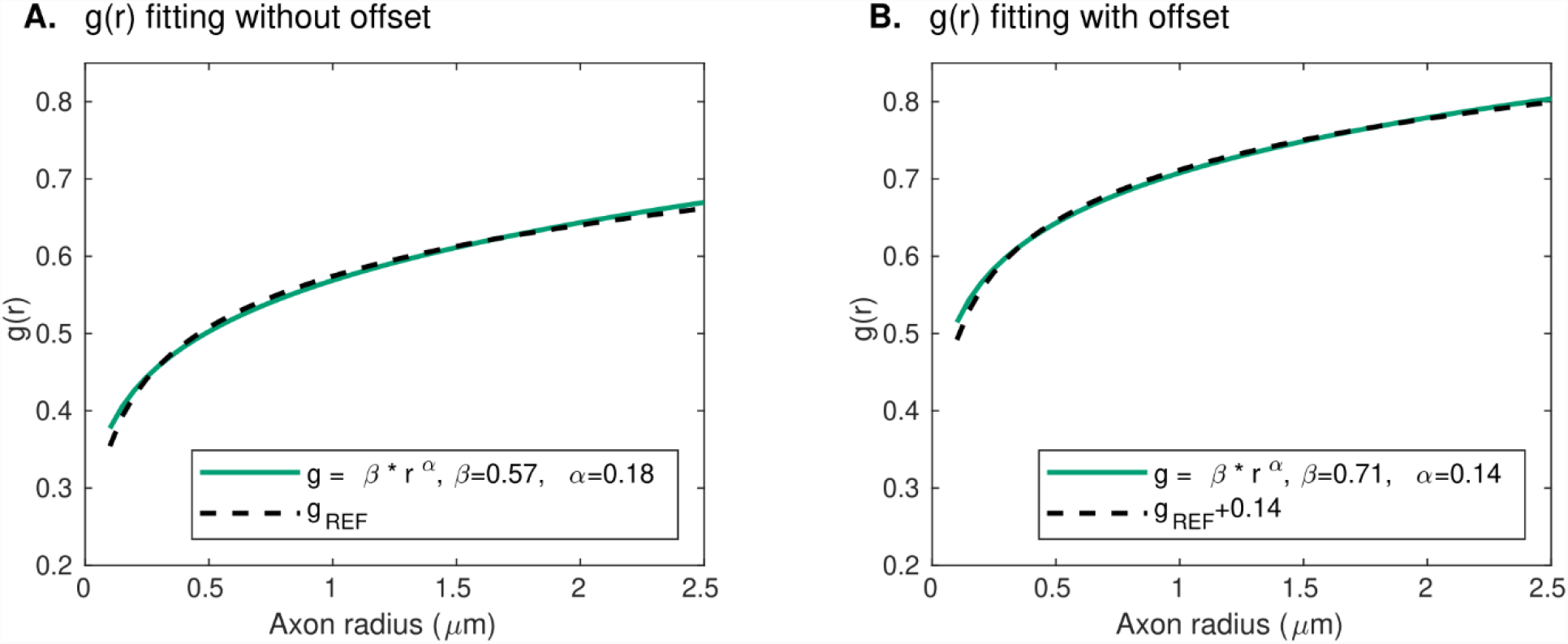
Radius dependence of the g-ratio. (A) An excellent agreement (R^2^=0.99) is found between the proposed power-law (green line) and reference histological measures of the g-ratio (*g*_*REF*_ (Ikeda and Oka, 2012), dashed line), with *α*=0.18 and *β*=0.57 *µm*^−*α*^. (B) The higher axonal g-ratios in the human central nervous system were accounted for by adding an offset of 0.14 to the reference peripherical nervous system data. The agreement between the proposed power-law and the reference *g*_*REF*_ remains very high (R^2^=0.99), with *α*=0.14 and *β*=0.71 *µm*^−*α*^.

### 2.2 Modeling of the in-vivo data

In agreement with West et al., 2016, the MRI g-ratio is written as an ensemble average of the axonal g-ratios within each image voxel, weighted by the axons’ cross-sectional area:

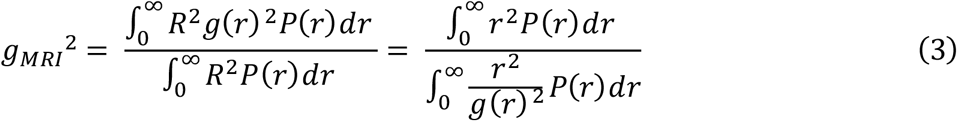

where *R* is the fiber radius (*R* = *r*/*g*(*r*)). From Eqs (1) and (2), Eq. (3) becomes (see Suppl. Appendix A):

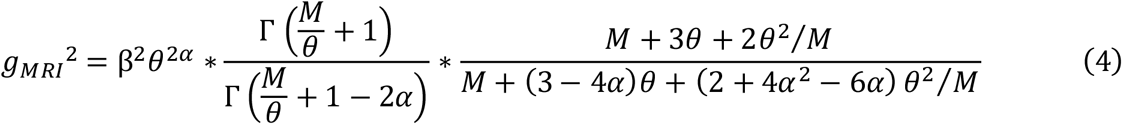

As described in Waxman and Bennett, 1972, axonal conduction velocity (*v*) can be derived from the morphological properties of axons using: 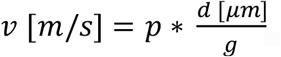, where *p* (∼5.5-6.0) represents the contribution of additional axonal factors to the propagation of action potentials (*e*.*g*. length of Ranvier nodes, electrical properties of the myelin membranes). Assuming an equal contribution from all axons to the conduction velocity *V* measured with EEG, we obtain:

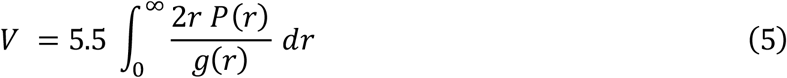

From Eqs (1) and (2), Eq. (5) becomes (see Suppl. Appendix A):

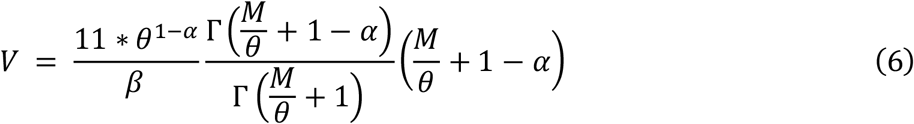

### 2.3 Model parameters

The proposed model (Eqs. (4) and (6)) includes 4 free parameters, pertaining to the distribution of axonal radius (*M* and *θ*) and to the radius-dependent axonal g-ratio (*α* and *β*). However, our proposed model uses only two data types acquired in-vivo (MRI g-ratio and EEG-based axonal conduction velocity). It is therefore necessary to set 2 parameters to reference values.

In general terms, the choice of model parameters to estimate should take into consideration the neural mechanisms in each possible application study of this model. The exponent *α* of the power-law *g*(*r*) represents the rate of change in thickness of the myelin sheath with axonal radius and may be a parameter of interest when considering changes in neuronal shape that differentially affect axons of different sizes. In contrast, the scale parameter *β* equally affects axons of all sizes. Concerning the distribution of axonal radius, we highlight that histological studies across white matter tracts and animal species have reported that, for reasons pertaining to brain size limitations and metabolism, the mode *M* of the axonal radius distribution remains largely constant (Liewald et al., 2014 and Tomasi et al., 2012). This motivates setting *M* to a constant value from the histological literature and estimating the tail parameter *θ* from the in-vivo data.

The choice of constant model parameters may also be guided by the impact of inaccuracies on the estimated morphological features. For instance, numerical simulations of the estimates computed from *g*_*MRI*_ and *V* values (using the procedure described in section 3.3) show that a 10% bias on the value of *α* leads to an average bias of ∼10% and ∼15% for *M* and *θ* respectively and a ∼0.5% bias on the estimates of *M* and *θ* for representative values of *g*_*MRI*_ and *V* (*g*_*MRI*_=0.72 and *V*=10 m/s, see section 4.1) (Suppl. Fig. 1A). Similarly, a 10% bias in *M* leads to an average bias of ∼1% and ∼22% for *β* and *θ* respectively and a bias <1% for *β* and ∼16% for *θ* for the representative values of *g*_*MRI*_ and *V* (Suppl. Fig. 1B). However, a 10% bias on the value of *β* leads to an average bias of ∼155% and ∼92% for *M* and *θ* respectively and a bias of ≥100% in the *M* and *θ* estimates for the same representative values of *g*_*MRI*_ and *V* (Suppl. Fig. 1C).

From the considerations above, in this study, we chose to set *α* to 0.14 (section 2.1) and *M* to 0.40 *µ*m (Tomasi et al., 2012; Liewald et al., 2014) and to estimate the parameters *β* and *θ*.

## 3 Materials and Methods

### 3.1 Numerical simulations

Numerical simulations were conducted using Eq. (4) and (6), to examine the values of the model parameters (*M, θ, α*, and *β*) obtained from combinations of the in-vivo data (*g*_*MRI*_ and *V*) using the procedure described in section 3.3, and vice versa.

In order to highlight the range of in-vivo data values compatible with the proposed model, estimates of *M* and *θ* were computed across a large range of *g*_*MRI*_ and *V* values, setting *α*=0.14 and *β*=0.71 (section 2.1). The range of in-vivo data values compatible with the proposed model was determined by comparison of the *M* and *θ* estimates against reference literature values.

Subsequent numerical simulations were conducted by estimating the parameters *β* and *θ* across the range of compatible in-vivo data values, setting *α*=0.14 and *M* to 0.40 *µ*m (section 2.3). In particular, we examined the range of the parameters *β* and *θ* across values of *g*_*MRI*_ and *V* reported in the literature. The variability of the model parameter estimates in the presence of noise was also investigated. Simulated estimates of the in-vivo data *g*_*MRI*_ and *V* were computed from combinations of *θ* and *β* values using Eq. (4) and (6). Noise was added to the computed *g*_*MRI*_ and *V* values with a standard deviation of 0.03 for *g*_*MRI*_ and 0.50 m/s for *V*, representative of intra-subject variability in in-vivo data. To replicate in-vivo conditions, 700 samples of *g*_*MRI*_ and one sample of *V* were taken from the resulting distributions of *gMRI* and *V*, and estimates of *θ* and *β* from noisy data were calculated. This process was repeated 2000 times and the standard deviation of the *θ* and *β* estimates across repetitions was computed as a measure of their variability.

Finally, we set out to investigate the effect of using a group averaged conduction velocity rather than subject-specific velocities. We selected 15 samples of *g*_*MRI*_ and *V* values from distributions with means of 0.70 and 10 m/s and standard deviations of 0.05 and 0.80 m/s respectively, representative of inter-subject variability in in-vivo data. From these simulated in-vivo data, reference values of *β* and *θ* were calculated. Estimates of *β* and *θ* were also computed from the average of the 15 samples of *V*. We then estimated the bias between the reference *β* and *θ* values and those obtained from the average value of *V*.

### 3.2 In-vivo data

We acquired data from 17 right-handed healthy volunteers. All participants had normal or corrected-to-normal vision and hearing and had no history of psychiatric or neurological disorder. Participant handedness was evaluated with the Edinburgh Handedness Inventory (Oldfield, 1971). All participants gave written informed consent and received 80 Swiss Francs as monetary compensation. The study was approved by the local ethics committee.

MRI data quality was assessed using the Motion Degradation Index (MDI) described in (Castella et al., 2018; Lutti et al., 2022). MDI values were ≲ 4 s^-1^ for the PD- and T1-weighted raw images, and ≲ 5 s^-1^ MT-weighted raw images, indicative of good quality images ((Lutti et al., 2022), see Suppl. Fig. 2). Three participants were excluded due to artifacted EEG recordings. The final sample consisted of 14 participants (6 females; age = 27.14±3.86 years). Three participants had left eye dominance, as determined by the eye viewing an object at a distance when the participant looks through a small opening (Miles, 1930). An overview of the data processing pipeline of the in-vivo data is shown in Fig. 1A.

#### 3.2.1 MRI-based estimation of the g-ratio

MRI data were collected on a whole-body 3T MRI system (Magnetom Prisma; Siemens Medical Systems, Erlangen, Germany) using a 64-channel receive head coil at the Laboratory for Neuroimaging Research, Lausanne University Hospital.

##### Structural MRI acquisition

A 3D structural T1-weighted Magnetization-Prepared Rapid Gradient-Echo (MPRAGE) image was acquired with a 1 mm^3^ isotropic voxel size and a matrix size of 176 × 232 × 256. TR/TE= 2000/2.39 ms. TI/α = 920 ms/9º. Parallel imaging (acceleration factor 2, GRAPPA reconstruction) was used along the phase-encoding direction (Griswold et al., 2002). The total acquisition time was 4 minutes.

##### Relaxometry MRI acquisition

The relaxometry MRI protocol consisted of multi-echo 3D fast low angle shot (FLASH) acquisitions with magnetization transfer-weighted (TR/α = 24.5 ms/6º, 6 echos), proton density-weighted (TR/α = 24.5 ms/6º, 8 echos) and T1-weighted (TR/α = 24.5 ms/21º, 8 echos) (Melie-Garcia et al., 2018) image contrasts (Fig. 1A). The echo spacing and minimal echo time were both 2.34 ms. The MR images had a 1 mm^3^ isotropic voxel size. Parallel imaging (acceleration factor 2, GRAPPA reconstruction) was used along the phase-encoding direction (Griswold et al., 2002) and Partial Fourier (acceleration factor 6/8) was used along the partition direction. B1-field mapping data was acquired to correct for RF transmit field inhomogeneities (Lutti et al., 2010, 2012): 4 mm^3^ voxel size, TR/TE = 500/39.1 ms. B0-field mapping data was acquired to correct for image distortions in the B1 mapping data: 2D double-echo FLASH, TR/ α = 1020 ms/90°, TE1/TE2 = 10/12.46 ms, BW = 260 Hz/pixel, slice thickness = 2 mm. The total acquisition time was 27 minutes.

##### Diffusion-weighted imaging (DWI)

Diffusion-weighted MRI data were acquired using a 2D echo-planar imaging sequence (TR/TE = 7420/69 ms) along 15, 30, and 60 diffusion directions with b = 650/1000/2000 s/mm^2^, respectively (Fig. 1A). 13 images with b = 0 were acquired, interleaved throughout the acquisition (Slater et al., 2019), making a total of 118 isotropically distributed directions. Images had a 2 mm^2^ isotropic voxel size and a matrix size of 96 × 106, with 70 axial slices. Parallel imaging was used along the phase-encoding direction (acceleration factor 2; GRAPPA reconstruction). The total acquisition time was 15 minutes.

##### Estimation of MRI quantitative maps

Maps of Magnetization Transfer (MT_sat_) were computed from the raw FLASH images as in (Helms et al., 2008a, 2008b) (Fig. 1A). The map computation was conducted using the hMRI toolbox (Tabelow et al., 2019) and included corrections for local RF transmit field inhomogeneities (Helms et al., 2008a) and for imperfect RF spoiling (Preibisch and Deichmann, 2009).

DWI data were corrected for geometrical distortions, using *eddy* from FMRIB’s Diffusion Toolbox (Andersson and Sotiropoulos, 2016), and for echo-planar imaging susceptibility distortions using the SPM12 fieldmap toolbox (Hutton et al., 2002). DWI images were aligned to the MT_sat_ map using SPM12 and a rigid body transformation. Finally, maps of the isotropic diffusion (*v*_*iso*_) and intracellular (*v*_*ic*_) compartments volume fractions were computed from the DWI data using the NODDI model (Zhang et al., 2012) and the AMICO toolbox (Daducci et al., 2015) (Fig. 1A).

Maps of the MRI g-ratio were estimated from: 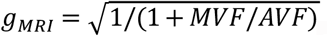, where MVF and AVF are the myelin and the axonal volume fractions, respectively (Stikov et al., 2015) (Fig. 1A). The MVF maps were estimated from the MT_sat_ maps according to: *MVF = αMT*_*sat*_, where the calibration factor *α* was set by assuming a median *g*_*MRI*_ value of 0.7 in the splenium of the CC of 11 subjects of a separate cohort (*α =* 0.23) (Campbell et al., 2018; Slater et al., 2019). The AVF maps were estimated as: *AVF =* (1 − *αMT*_*sat*_)(1 − *v*_*iso*_)*v*_*ic*_ (Stikov et al., 2015).

The Freesurfer 6.0 software (Schiffler et al., 2017) was used to delineate each subject’s Brodmann areas 17 and 18 (Fischl et al., 2008) from their MPRAGE image, corresponding to the primary (V1) and secondary (V2) visual cortical areas, respectively. These regions of interest (ROIs), henceforth called ‘V1V2’, were grouped and registered to the diffusion data using FMRIB’s Linear Image Registration Tool (Jenkinson and Smith, 2001).

Tractography analysis was conducted using the mrTrix software (Tournier et al., 2019): whole-brain anatomically constrained tractography was performed using the iFOD2 algorithm, followed by the SIFT2 to penalize streamlines with a reduced agreement with diffusion data. The visual transcallosal tract was isolated by selecting the streamlines connecting the V1V2 ROIs in each hemisphere across the CC. This tract was used to extract samples from the MRI g-ratio maps (Fig. 1A).

#### 3.2.2 EEG-based estimation of the IHTT

##### Experimental paradigm

We implemented the visual Poffenberger paradigm on a 61 cm widescreen (60 Hz refreshing rate), in line with the literature (Westerhausen et al., 2006; Whitford et al., 2011; Friedrich et al., 2017; Chaumillon et al., 2018). The experiment was administered using Psychtoolbox-3.0.16 in Matlab (R2019b, The Mathworks, Natick, MA). Participants were comfortably seated on a chair in a dimly lit room, at a standardized distance of 80 cm from the screen, thus 1 cm on the screen represented 0.72° of the visual angle. Each trial consisted of the presentation of a black and white circular checkerboard with a pattern reversal of 15 Hz on a grey background (20 cd/m^2^) and with a duration of 100 ms. The stimuli were of 4º diameter and their outer edge appeared at 6° horizontal and 6° vertical distance from the centrally-located fixation cross (0.8º size) to the lower left or right visual half-field. The acquisition was structured in 6 blocks with an approximate duration of 6 minutes each; a short break was allowed between each experimental block. Each block consisted of 205 trials (95 for right visual field, 95 for left visual field, and 15 where no stimulus appeared), presented in a pseudorandom order and with inter-trial intervals randomly assigned between 1.0 and 2.0 s. Participants were instructed to avoid unnecessary movements and to press a button as quickly as possible after the appearance of a stimulus while keeping their gaze on the fixation cross. Responses were given with the index finger via a keyboard button press placed centrally to the subject’s body. The administered blocks alternated between left and right-hand index finger button presses (3 blocks per hand).

##### EEG data acquisition

Continuous 128-channel EEG was recorded using the Micromed recording system (Micromed SystemPlus Evolution, Mogliano Veneto, Italy) and an Ag/AgCl electrode cap (waveguard™original, ANT Neuro, Hengelo, Netherlands) at a sampling rate of 1024 Hz with FPz as the reference electrode and AFFz as the ground electrode. Two additional horizontal EOG electrodes were attached to the outer canthi of each eye. Electrode impedance was kept below 20 kΩ. Electrode positions and head shape were acquired for each participant using the xensor™ digitizer (ANT Neuro, Hengelo, Netherlands).

##### EEG data analysis

EEG data analysis utilized custom-made MATLAB (R2021a, The Mathworks, Natick, MA) scripts and open-source toolboxes Fieldtrip (version 20191206, (Oostenveld et al., 2011), EEGLAB (version 13.4.4b, (Delorme and Makeig, 2004)) and Brainstorm (Tadel et al., 2011). Continuous raw EEG was bandpass-filtered between 0.1 and 40 Hz (digital filters). EEG epochs were extracted from the filtered data ranging from −100 to 300 ms relative to visual stimulus onset. Artifact trials were removed based first, on visual inspection. Next, we identified and removed components containing eye movement related artifacts by running an Independent Component Analysis based on the *runica* algorithm (Bell and Sejnowski, 1995). Epochs containing additional artifacts were identified based on a threshold of 80 *µ*V and excluded from further analysis. Across participants, an average of 20.5% (SD: 10.1%, range: 43 to 220 trials) and 21.3% (SD: 11.0%, range: 42 to 206 trials) of the trials were rejected for the left and right visual field stimulation, respectively. Artifact electrodes were identified based on a threshold of 80 *µ*V and were interpolated using the nearest neighbors. On average, 5.9% of electrodes (SD: 2.4%, range 3 to 14 electrodes) were interpolated across participants. Epoched data were re-referenced to the average reference. We removed DC drift by subtracting the average within each epoch.

Source reconstruction was performed to identify the neural origins underlying the visual evoked response to the left and right hemifield visual stimuli (Fig. 1A). Virtual sensors from artifact-free EEG data were calculated using the minimum-norm current density method (Hämäläinen, 2010) as implemented in Brainstorm. The MRI image of each subject was registered to the electrode positions using an iterative algorithm that finds the best fit between the head shape obtained using the MRI data and that obtained via EEG digitization. Surface reconstructions were obtained using a 3-layer Boundary Element Method (Kybic et al., 2005; Gramfort et al., 2010) model on each subject MRI image. The source grid was defined with 15000 points on the grey matter. This way, we estimated the current densities (CD, pA.m) for each condition, source, and time point within each subject. The CDs were extracted for the V1V2 ROI defined in section 3.2.1 – for the left and right brain hemisphere – for each trial and each subject. Next, V1V2 ROI CDs were averaged across trials within each subject. Given that cortical anatomies vary considerably across participants due to the folding patterns of each individual, current source density maps have ambiguous signs on the group level. Consequently, we took the absolute value of the CDs of each subject before computing a group average.

##### Estimation of the IHTT and conduction velocity

We assumed that lateralized visual stimuli would induce first, a contralateral activation of the visual cortex, followed by an activation of the ipsilateral cortex. This visual information transfer is assumed to be achieved through the CC (Marzi, 1999).

For the IHTT estimation, we identified the first peaks of activation in each hemisphere based on the maximum of the average current density value at the group level (Fig. 1A). IHTT was calculated as the latency difference between the ipsilateral and contralateral activation peaks on the group average CDs within the V1V2 ROI. A Wilcoxon signed-rank test (p<0.05; *signrank*, in MATLAB) comparing the CDs across participants at these two maxima to the time-average baseline values was used to evaluate the significance of these activations as evoked activity in response to the visual stimuli.

To obtain a confidence interval on the computed IHTT, we subdivided the artifact-free EEG trials available for each participant into four non-overlapping splits. We then repeated four times the CD estimation at group-level where each subject contributed with data coming from one of the split in order to obtain four independent estimation of the IHTT. This allowed us to obtain a standard deviation on the IHTT estimation, which we used to define a confidence interval on the IHTT estimation: *[IHTT-standard deviation; IHTT+standard deviation]*.

Of note, we assumed that the right eye dominance of the majority of the included participants elicited a more reliable estimation of the evoked activity following the left visual stimuli in comparison to the right. For this reason, here, we used the IHTT estimation following the left hemifield visual stimuli.

At the individual level, we extracted the visual transcallosal tract length as the tractography-based mean streamline length. *V* was calculated by dividing the tract length by the IHTT.

### 3.3 Estimation of axon morphology from in-vivo data

Estimation of axonal morphological features from the in-vivo MRI and EEG data was implemented using MATLAB-based custom-made analysis scripts. The MRI g-ratio samples along the visual transcallosal tract and the estimate of the IHTT were used to estimate model parameter values using Eq. (4) and (6) (see section 2.1, Fig. 1B). This was achieved using MATLAB’s nonlinear least-square routine (*lsqnonl*) with a trust-region-reflective minimization, which minimizes the sum of the squares of the residuals. The initial conditions used to ensure convergence of the fitting routine were set to *β*=0.70 *µm*^−*α*^ and *θ*=0.10 *µ*m. All codes will be made available to general public upon publication of this manuscript.

## 4 Results

### 4.1 Numerical Simulations

We investigated the range of in-vivo data compatible with the proposed model by considering the values of the model parameters *M* and *θ* computed from each combination of *g*_*MRI*_ and *V* (Fig. 3). Combinations of large values of *g*_*MRI*_ and low values of *V* lead to unrealistically low values of *M* (∼10^− 14^ *µ*m). Conversely, small values of *g*_*MRI*_ and large values of *V* lead to excessively large values of *M* (∼1 *µ*m) and low values of *θ* (∼0.001 *µ*m) (Aboitiz et al., 1992; Caminiti et al., 2009, 2013; Tomasi et al., 2012; Liewald et al., 2014).

**Figure 3.**
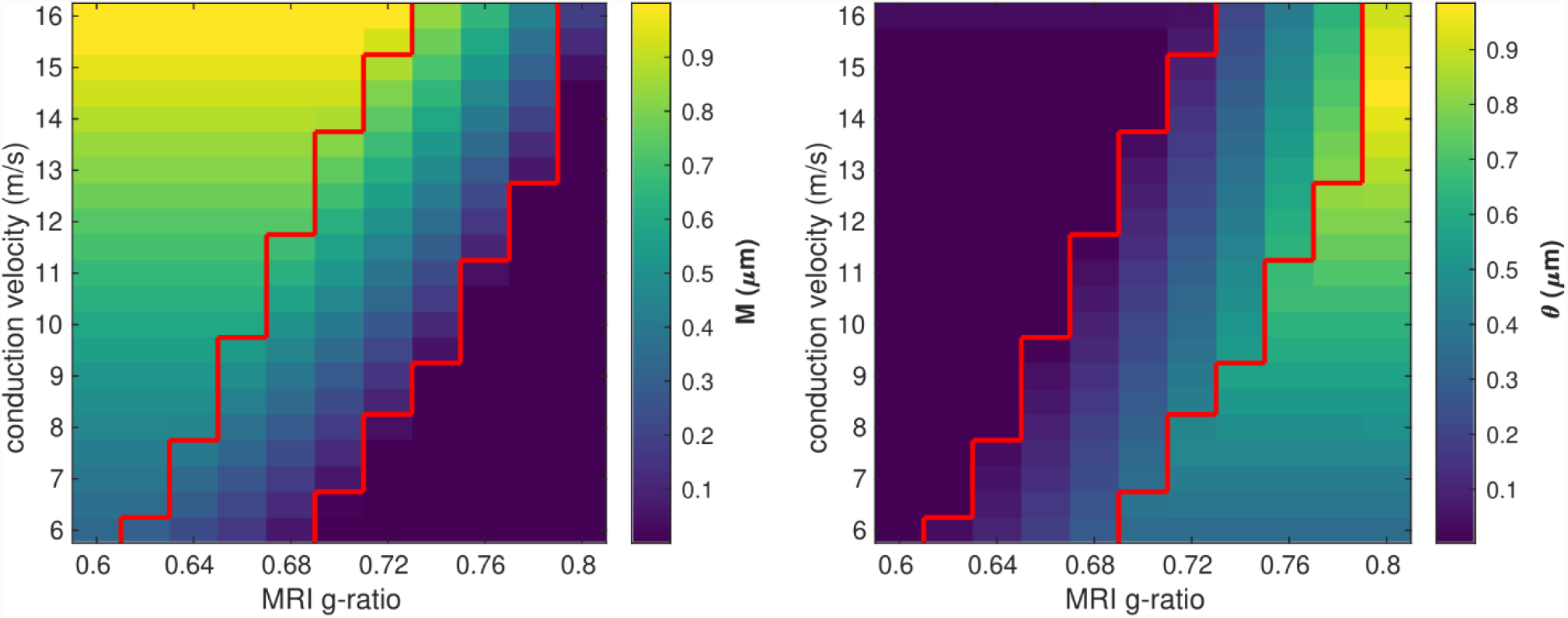
Range of in-vivo data compatible with the proposed model. Combinations of *g*_*MRI*_ and *V* that lead to biologically plausible values of the model parameters (0.05<*M*<0.9 *µ*m and 0.01< *θ*<0.9 *µ*m) are located between the two red contour lines.

Fig. 4 shows the range of the model parameters *θ* and *β* across a large range of *g*_*MRI*_ and *V* values. According to the literature, frontal transcallosal white matter exhibits MRI g-ratio values of ∼0.62 (Mohammadi et al., 2015; Slater et al., 2019) and conduction velocities of ∼8 m/s (Caminiti et al., 2013) (see red cross in Fig. 4A). The proposed model yields *θ*∼0.05 *µ*m for such combination of *g*_*MRI*_ and *V*, equivalent to a mean axonal radius of 0.45 *µ*m (Fig. 4B). This value is consistent with histological analyses of such white matter fibers, with a narrow range of axonal radius (mean∼0.48 *µ*m, Caminiti et al., 2009). For visual transcallosal white matter (MRI g-ratio ∼0.72; conduction velocity ∼10m/s (Caminiti et al., 2013; Mohammadi et al., 2015; Slater et al., 2019)), the proposed model yields *θ*∼0.23 *µ*m (see black cross in Fig. 4A), leading to a mean axonal radius of ∼0.63 *µ*m (Fig. 4B). This value is consistent with histological analyses of such white matter fibers, with a broad range of axonal radius (mean∼0.62 *µ*m, Caminiti et al., 2009). For both types of white matter tract, fibers with a radius above 1.5 *µ*m represent less than 4% of the total number of fibers, consistently with previous histological studies (Aboitiz et al., 1992; Caminiti et al., 2009; Liewald et al., 2014). The value of the model parameter *β* for frontal and visual transcallosal white matter were 0.68 and 0.73 *µm*^−*α*^ respectively (Fig. 4C and 4D), in line with histological analyses of the axonal g-ratio in the genu and splenium of the CC (Stikov et al., 2015).

**Figure 4.**
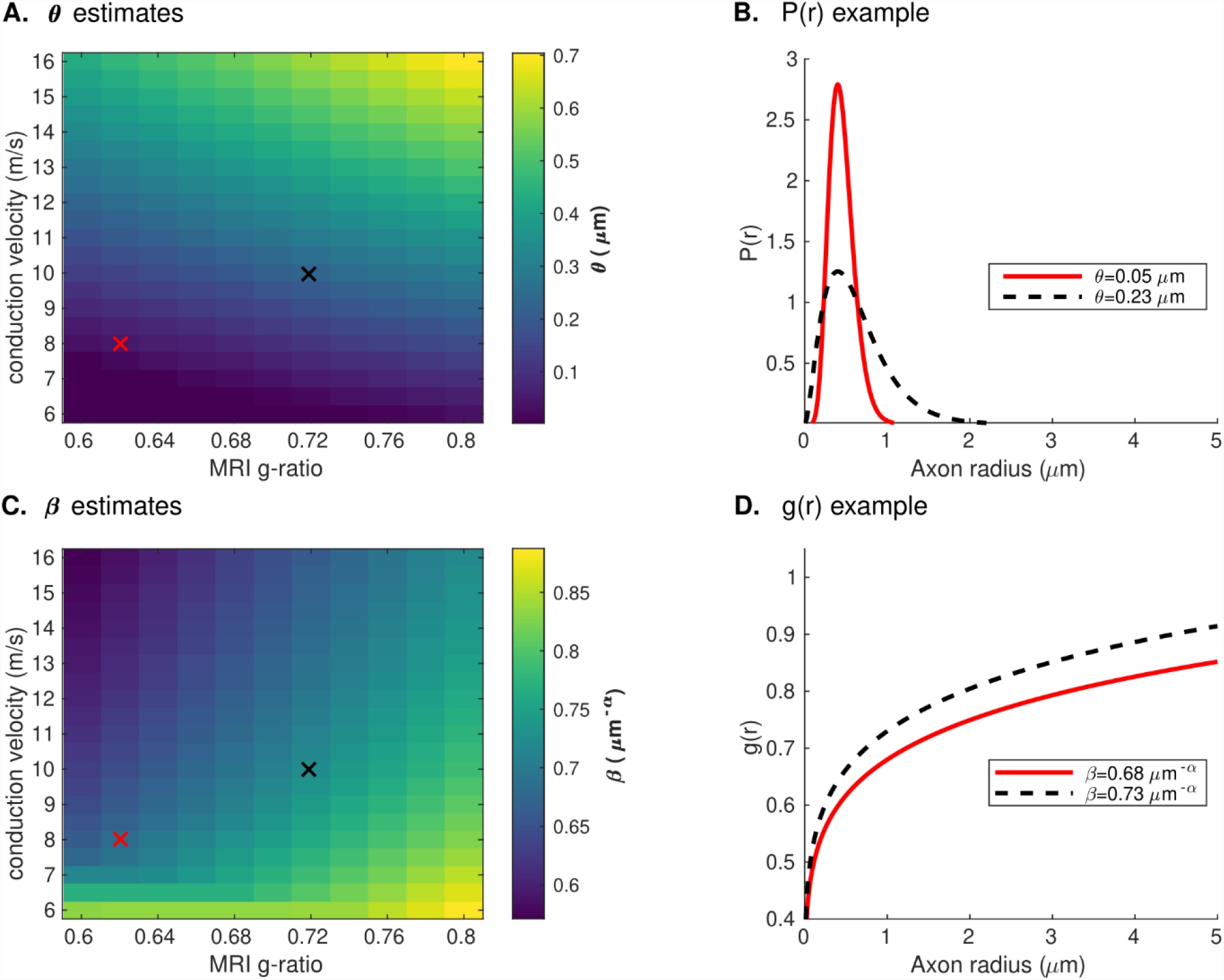
Range of the model parameters θ and β. (A) Range of the model parameter θ, computed from combinations of simulated in-vivo g_MRI_ and V. (B) Axonal radius distributions representative of frontal and visual transcallosal white matter (θ =0.05 and 0.23 μm). (C) Range of the model parameter β, computed from combinations of simulated in-vivo g_MRI_ and V. (D) Dependence of the fiber g-ratio on the axonal radius, representative of frontal and visual transcallosal white matter (β=0.68 and 0.73 μm^−α^). In (A) and (C), the red and black crosses illustrate values of g_MRI_ and V for frontal and visual transcallosal tracts, taken from the literature (Caminiti et al., 2013; Mohammadi et al., 2015; Slater et al., 2019).

Fig. 5 shows the variability of the *θ* and *β* estimates in the presence of noise in the in-vivo data, computed from combinations of conduction velocity and MRI g-ratio across a plausible range with 700 *g*_*MRI*_ samples and one *V* sample. The highest errors in *θ* (∼35%) are found for the smallest values of *θ*, with only a small effect of the parameter *β*. The highest errors in *β* (∼1%) are obtained from a combination of small values of *θ* and large values of *β*. Suppl. Fig 3 shows the dependence of the variability of the *θ* and *β* estimates on the number of samples of *V* and *g*_*MRI*_ (*β*=0.71 *µm*^*− α*^ and *θ*=0.22 *µ*m). With only one estimate of *V*, as was the case in this study, errors of up to 11% on the *θ* estimates are observed. The errors on the *β* estimates are mostly driven by the number of *g*_*MRI*_ samples included in the estimation: for 100 *g*_*MRI*_ samples or more, these errors are below 1% regardless of the number of *V* samples.

**Figure 5.**
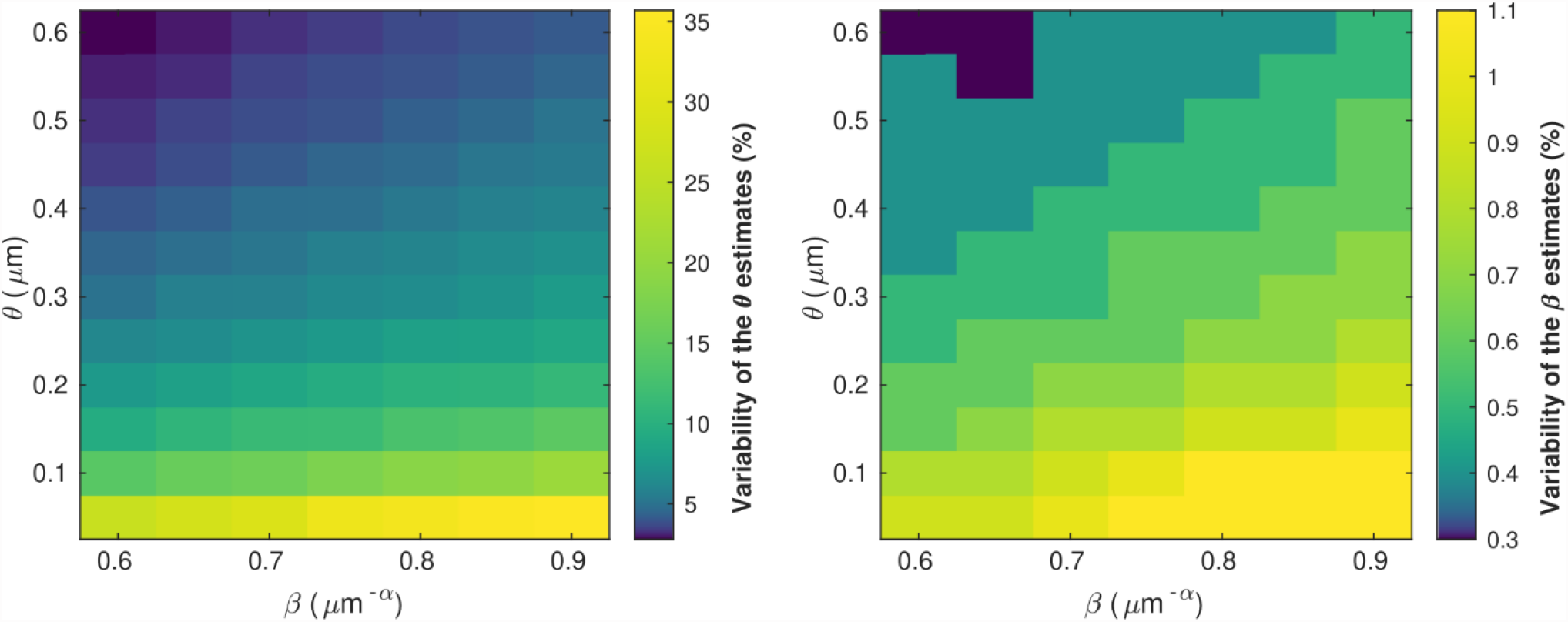
Variability of the θ (left) and β (right) parameter estimates due to noise in the in-vivo samples of g_MRI_ and V. The highest errors in θ (∼35%) are found for the smallest values of θ, with only a small effect of the parameter β. The highest errors in β (∼1%) are obtained from a combination of small values of θ and large values of β.

Fig. 6 shows the bias in the *θ* and *β* estimates arising from an estimation of conduction velocity from a cohort of participants. The bias in *θ* and *β* is on average 12% and 0.9% across participants, reaching up to 40% and 3%, respectively.

**Figure 6.**
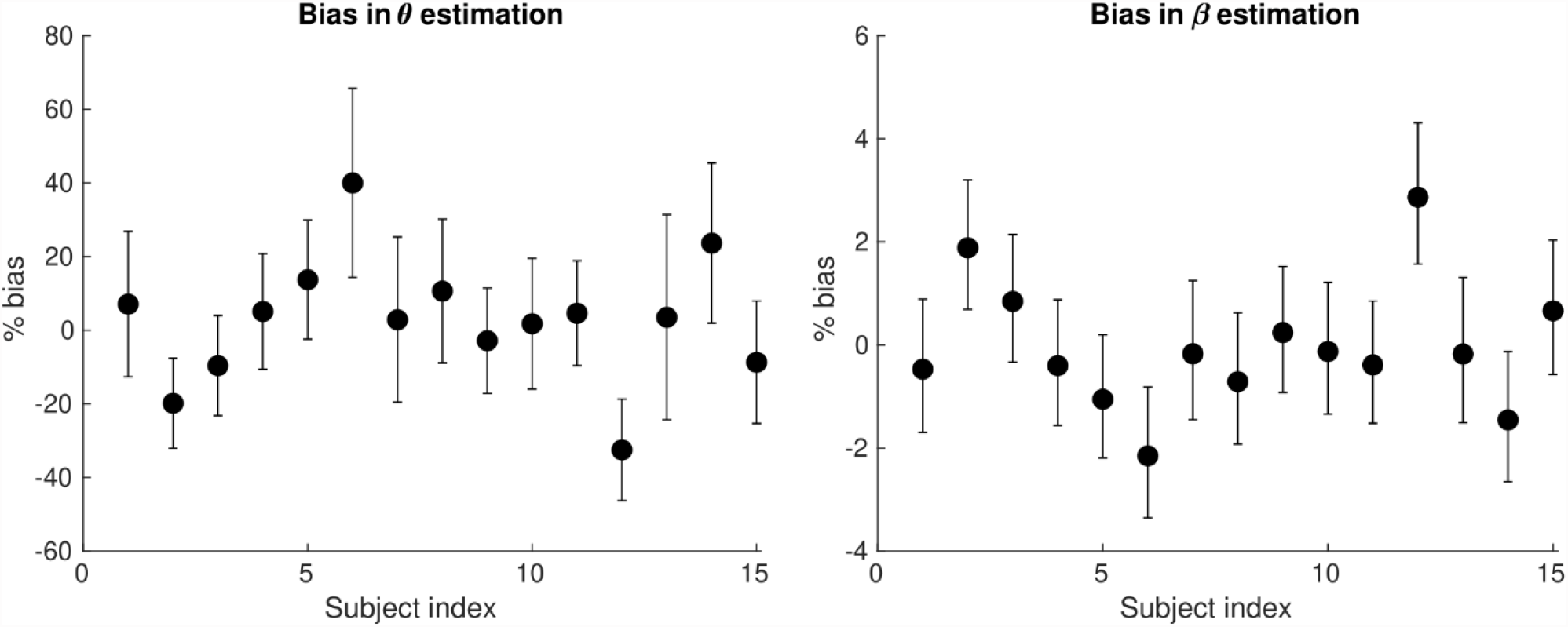
Bias of the *θ* (left) and the *β* (right) parameter estimates arising from the computation of axonal conduction velocity across a group of subjects (N=15). The error bars indicate the standard deviation on the parameter estimates. The bias of *θ* and *β* reach up to 40% and 3% respectively.

### 4.2 Estimation of the IHTT and conduction velocity

At the sensor level, the group-averaged visual evoked response revealed a positive peak activation, between 115 ms and 134 ms post-stimulus onset, contralaterally to the stimulus presentation (Fig. 7, top). The latency of this positive activation corresponds to the expected latency of the P100 component (Di Russo et al., 2001) and was followed by asymmetrically distributed voltage topographies with maximal voltage values at posterior sites at latencies starting at approximately 153 ms post-stimulus onset (Fig. 7, bottom). After approximately 180 ms, post-stimulus onset voltage topographies showed an ipsilateral positivity to the stimulus presentation, suggesting that the P100 activation has traveled to the opposite hemisphere.

**Figure 7.**
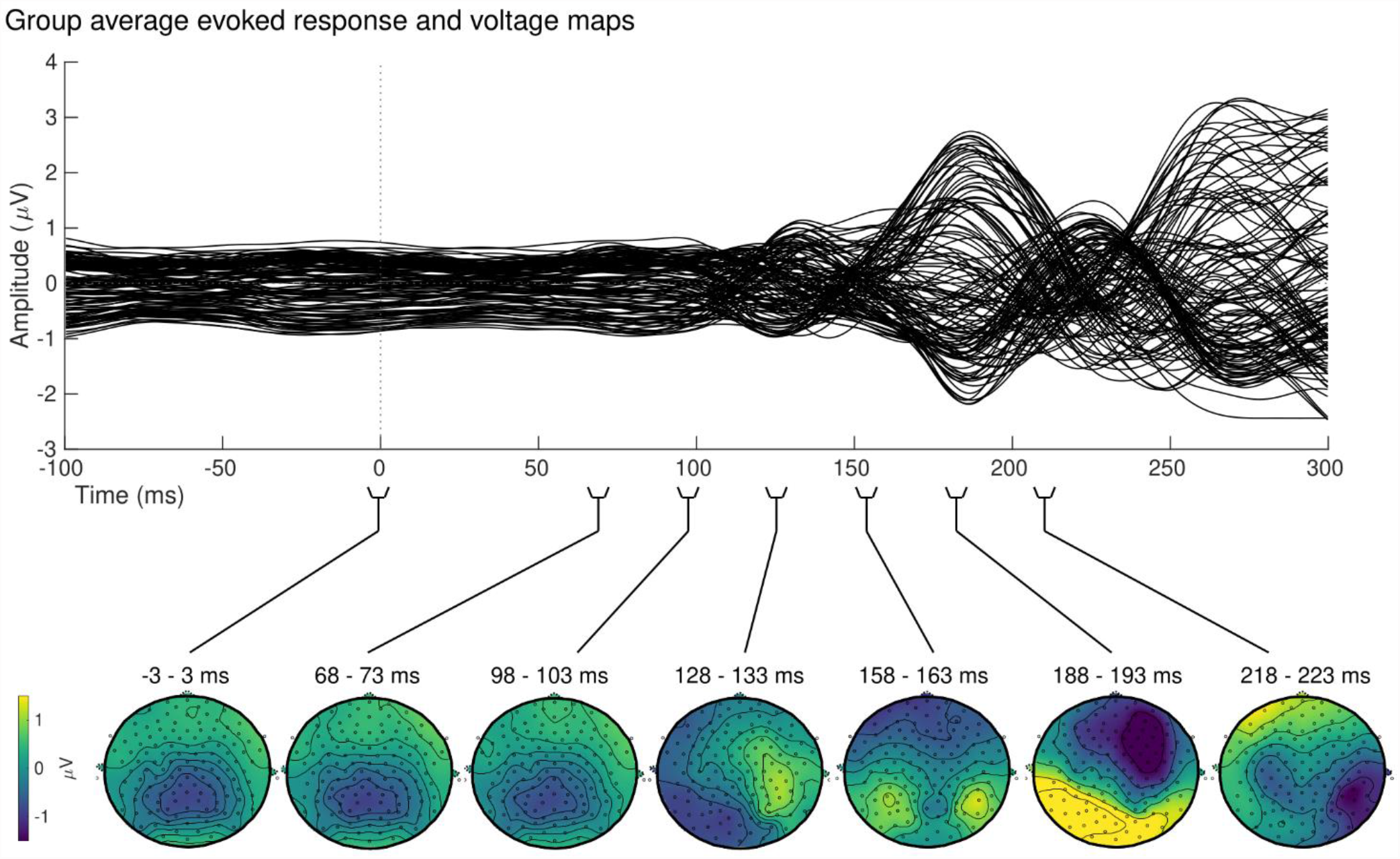
Group sensor space results. Grand-average evoked response (top) and corresponding voltage topographic maps (bottom) after left visual field stimulation. Topographic maps are shown on a flattened electrode layout with anterior regions at the top and posterior regions at the bottom. Time 0 ms identifies the onset of the stimulus presentation. Positive amplitudes (yellow) were observed in the hemisphere contralateral to the stimulus stimulation between 115 and 134 ms, followed by a bilateral positivity starting at approximately 153 ms.

Qualitative observations at the electrodes-level were confirmed by the source reconstruction results. Group-averaged absolute CD exhibited a sharp increase at approximately 100 ms post-stimulus onset in the right hemisphere compared to the baseline, peaking at 141ms (Fig. 8A and Fig 8B for an overview of the spatial distribution of the CD for an exemplar subject). The group-averaged absolute CD on the left visual hemisphere followed the right visual hemisphere, peaking later on at 152ms.

**Figure 8.**
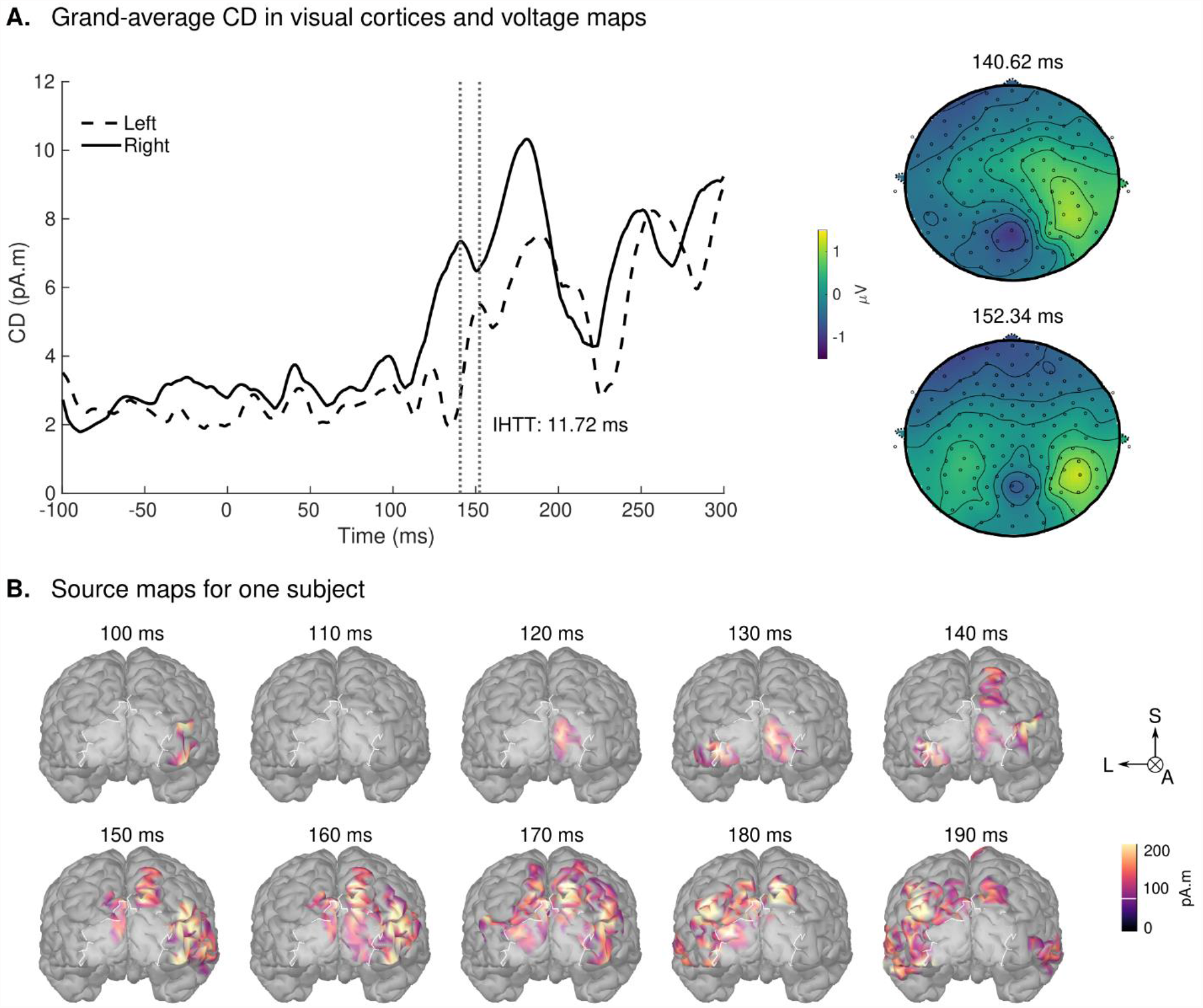
Source space results. (A) Group averaged CD for the left visual field stimulation (left panel). The mean activation in the V1V2 ROI on the left and right hemispheres are shown in dashed and solid lines, respectively. Time 0 ms identifies the onset of the stimulus presentation. A first peak on the right hemisphere is observed between 100 and 145 ms, followed by a peak on the left hemisphere after 145 ms. Vertical lines identify the peaks of activation in both hemispheres. IHTT was calculated as the difference between the peak of activation on the hemisphere ipsilateral (left) and contralateral (right) to the stimulation visual field: 11.72 ms. The grand-average sensor level topographic voltage maps corresponding to the identified peaks are shown on the right panel. (B) Spatial distribution of the CDs of one exemplar subject projected on this subject’s cortex for the left visual field stimulation during the post-stimulus period. Highlighted in white is the V1V2 ROI of interest. For the sake of clarity, we show only groups of source values that contained more than 30 vertices and were 36% above minimal activation. L: Left; S: Superior; A: Anterior.

The IHTT group estimation yielded a value of 11.72 ms with a standard deviation of 2.87 ms. The identified peaks were statistically significantly larger upon visual stimulation when compared to the baseline with p-values of p<0.01.

Using the estimated group IHTT, we calculated the conduction velocity for the visual transcallosal tract using each participant’s tract length. The average conduction velocity was 13.22±1.18 m/s across participants (Table 1).

**Table 1.** Conduction velocity estimates and corresponding confidence intervals for each subject in the visual transcallosal tract.

| Tract length<br>(mm) | Velocity<br>(m/s) | Velocity<br>confidence<br>interval (m/s) |
| --- | --- | --- |
| 155.03 | 13.23 | [10.63 ; 17.52] |
| 149.38 | 12.75 | [10.20 ; 16.88] |
| 133.43 | 11.38 | [9.15 ; 15.08] |
| 136.25 | 11.63 | [9.34 ; 15.40] |
| 154.32 | 13.17 | [10.58 ; 17.44] |
| 171.41 | 14.63 | [11.75 ; 19.37] |
| 157.94 | 13.48 | [10.83 ; 17.85] |
| 149.95 | 12.79 | [10.28 ; 16.94] |
| 152.98 | 13.05 | [10.49 ; 17.29] |
| 142.29 | 12.14 | [9.75 ; 16.08] |
| 155.49 | 13.27 | [10.66 ; 17.57] |
| 172.32 | 14.70 | [11.81 ; 19.47] |
| 184.48 | 15.74 | [12.64 ; 20.85] |
| 154.40 | 13.17 | [10.58 ; 17.45] |

### 4.3 Estimation of axonal morphology from in-vivo data

Estimates of the model parameters *θ* and *β* were computed using the proposed model, from the in-vivo samples of the MRI g-ratio along the visual transcallosal tract and estimates of conduction velocity. The average value of *θ* was 0.40±0.07 *µ*m across all subjects and ranged between 0.31 and 0.54 *µ*m (Fig. 9A). A *θ* value of 0.40 *µ*m is equivalent to a mean axonal radius of 0.80 *µ*m, in-line with previous estimates from histological studies (0.62 μm, Caminiti et al., 2009). For such a value of *θ*, axons with a radius above 2 *µ*m represent only <5% of the total fiber count, in agreement with histological studies that shows that axonal radius does not exceed ∼1.5–3 *µ*m in the human brain (Aboitiz et al., 1992; Caminiti et al., 2009; Liewald et al., 2014). The average value of *β* was 0.67±0.02 *µm*^−*α*^ across all subjects, and ranged between 0.64 and 0.70 *µm*^−*α*^ (Fig. 9B). Fig. 9C shows the axonal radius distribution *p(r)*for a representative subject (*θ* = 0.43 *µ*m), with a confidence interval of 0.27-0.69 *µ*m obtained from the IHTT estimation. The estimated value of *β* for this subject was 0.68 *µm*^−*α*^, with a confidence interval between 0.64 and 0.71 *µm*^−*α*^ (Fig. 9D).

**Figure 9.**
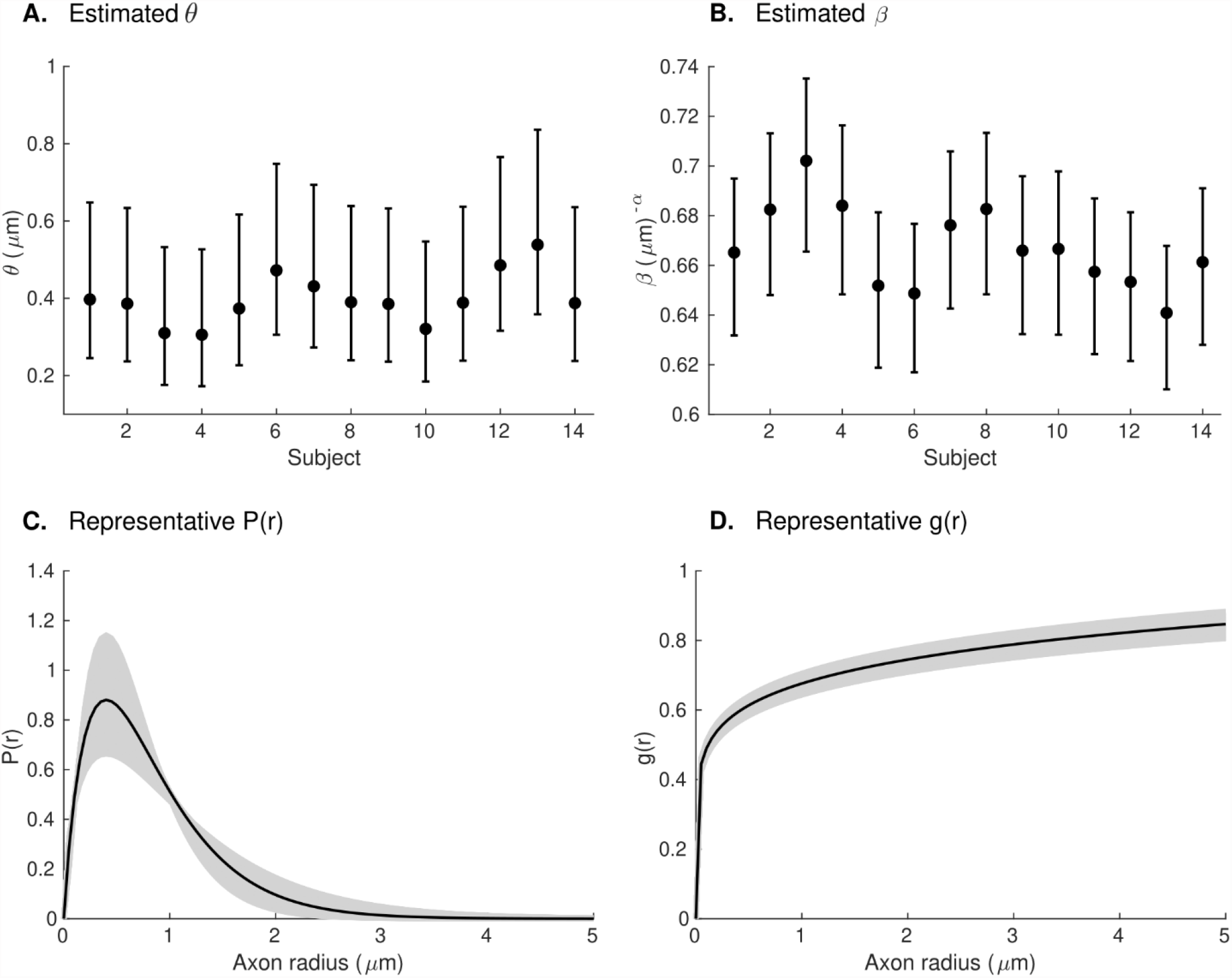
Estimates of axonal morphology obtained from the in-vivo *g*_*MRI*_ and IHTT samples. (A) Estimates of the model parameter *θ* – the width of the right tail of the axonal radius distribution – in the visual transcallosal tract across all study participants. (B) Estimates of the model parameter *β* – the scaling factor of the axonal g-ratio – in the visual transcallosal tract across all study participants. The error bars indicate the confidence intervals on the parameter estimates. (C) Exemplar axonal radius distribution in the visual transcallosal tract for a representative subject (*θ*=0.43 *µ*m; confidence interval: 0.27-0.69 *µ*m, shaded area). (D) Exemplar dependence of the fiber g-ratio on the axonal radius for the same subject (*β*=0.68 *µm*^−*α*^, confidence interval: 0.64-0.71 *µm*^−*α*^, shaded area).

## 5 Discussion

In this paper, we propose a novel method that allows the non-invasive estimation of morphological properties of white matter axons from in-vivo human data. This approach requires MRI g-ratio and EEG-based measures of axonal conduction velocity computed from estimates of the IHTT. From these measures, we estimated the axonal radius and myelination of axonal fibers, distinct histological features of white matter. These morphological features were assessed across the distribution of axons in the visual transcallosal tract, providing a detailed insight into the microscopic properties of these white matter fibers.

### 5.1 Estimation of axonal morphology from in-vivo data

The proposed model is based on an explicit link between the data acquired in-vivo and a limited set of histological properties of white matter axons. The MRI g-ratio is expressed as a function of the axonal radius distribution (*P*(*r*)) and the g-ratio of axonal fibers (*g*(*r*)), in-line with recent studies conducted using MRI and histology data (Stikov et al., 2011, 2015; West et al., 2016). Similarly, axonal conduction velocity – computed from the IHTT estimates – is an ensemble average across the same distribution *P*(*r*), assuming an equal contribution of all axons to the EEG data. As a result, both types of in-vivo data depend on the same properties of axonal fibers: *P*(*r*) and *g*(*r*). This approach is supported by recent findings which show that from the numerous histological determinants of conduction velocity (*e*.*g*. axonal radius, g-ratio, the conductance of ion channels, diameter and length of Ranvier nodes and internodes), the properties of axons that bring the largest contribution to the determination of conduction velocity are measurable with MRI (Drakesmith et al., 2019).

The mathematical definitions of *P*(*r*) and *g*(*r*) are grounded on well-established histological findings. *P*(*r*) is assumed to follow a gamma distribution, as commonly posited by models of axonal radius distribution (Assaf et al., 2008; Sepehrband et al., 2016). *g*(*r*), expressed using a power law, shows a high level of agreement with histological studies (Fig. 2, Ikeda and Oka, 2012; Gibson et al., 2014). Besides supporting the proposed model, the histological basis for the expressions of *P*(*r*) and *g*(*r*) allows freedom in the choice of the model parameters estimated from the data, because they reflect different properties of axon populations that can be set constant or variable according to their relevance in a neuroscience application of this model.

From the set of 4 model parameters, we opted to set the mode *M* of the axonal radius distribution to a constant value, based on the histological literature (Tomasi et al., 2012; Liewald et al., 2014). Similar ly, the parameter *α* was set from a calibration with histological findings (*g*_*REF*_) (Ikeda and Oka, 2012) (see section 2.1). The estimated parameters – *θ*, the right tail of *P*(*r*) and *β*, the scaling parameter of *g*(*r*) – enabled the simultaneous assessment of axonal radius and myelination, across the distribution of axons in the visual transcallosal white matter tract. The average value of *θ* was 0.40 *µ*m across participants, leading to a mean axonal radius of 0.80 *µ*m, in-line with previous estimates from histological studies (0.62 *µ*m, Caminiti et al., 2009). The average value of *β* was 0.67 across participants, consistent with histological measures of the axonal g-ratio in the splenium of the CC (Jung et al., 2018).

### 5.2 Estimation of the IHTT and conduction velocity

The IHTT was estimated from the group averaged CDs, where we could easily pinpoint the first two maxima of activation in the two hemispheres (Fig 8). The IHTT derived as a result of the time-interval separating the peak of CDs at the group level (11.72±2.87 ms) falls within the range (*i*.*e*. 8 and 30 ms) of previous IHTT estimates based a priori selection of voltage measurement at electrodes at occipital sites (*e*.*g*. Chaumillon et al., 2018; Saron and Foxe, 2003; Westerhausen et al., 2006; Whitford et al., 2011; Friedrich et al., 2017). In our study, we opted for a CD-based estimation of the IHTT as the closest reflection of the evoked neural activity in the regions of interest, consistent with the corresponding white matter tract selection (section 3.2.1). In addition, this approach helps to overcome the ambiguity of electrode selection in electrode-based IHTT estimations. Of high relevance, the conduction velocities we calculated are in agreement with the reported values by Caminiti and colleagues (Caminiti et al., 2013), wherein velocities of 10 m/s were obtained using histology samples in humans in the same fiber tracts we investigated. Our results are not only in-line with ex-vivo and IHTT studies but are also consistent with a more recent report that uses a different approach based on Bayesian networks to map the flow of information following left visual stimulation (Deslauriers-Gauthier et al., 2019). The authors observed a transfer of information from the right to the left occipital cortex, between 140 ms and 160 ms (Deslauriers-Gauthier et al., 2019), in agreement with the latency of the right and left activations observed in our analysis.

Ideally, IHTT estimates should be performed on the single-subject level to provide microstructure measures that are specific to each subject. However, identifying the ipsilateral peak of activation was not feasible in our entire participant population which prevented us from defining a robust criterion to estimate the IHTT. The reasons for this might be related to differences in brain morphology (Saron and Foxe, 2003), the number of available trials (Iacoboni and Zaidel, 2000), and/or ambiguity in the selection of the peak. In the literature, IHTT values have also been shown to be inconsistent across participants and counterintuitively, in some cases have even been calculated as negative (*e*.*g*. Saron and Davidson, 1989; Marzi et al., 1991; Westerhausen et al., 2006; Friedrich et al., 2017), the latter suggesting that interhemispheric transfer evaluated at the electrode and at the single-subject level can produce either inaccurate results or in disagreement with our assumptions that evoked activity produces a contralateral activation prior to the interhemispheric transfer.

Furthermore, our results could be improved with further development in the reliability of the inverse solution (Plomp et al., 2010; Mahjoory et al., 2017), which may allow a single-subject estimation of the IHTT.

### 5.3 Anatomical substrate for the interhemispheric transfer

The proposed model is based on the combination of a structural measure of white matter (MRI g-ratio) and a measure of brain function (axonal conduction velocity). The validity of this model relies on the assumption that both measures may be obtained for a given white matter tract. Anatomical delineation of a white matter tract is generally a routine procedure thanks to MRI tractography techniques (Caminiti et al., 2013; Horowitz et al., 2015; Tournier et al., 2019), allowing the sampling of the MRI g-ratio data along this tract (Schiavi et al., 2022). On the other hand, the underlying mechanisms and anatomy of the inter-hemispheric transfer of the evoked visual activity in Poffenberger’s paradigm are still under investigation. Previous literature has used this paradigm to demonstrated an increase in IHTT in acallosal subjects, highlighting the primary role of the CC in visual interhemispheric transfer (Marzi et al., 1991; Di Stefano et al., 1992; Aglioti et al., 1993; Tassinari et al., 1994). In addition, Westerhausen et al., 2006 showed that IHTT is significantly correlated with the structural integrity of the posterior CC, suggesting splenium fibers as the most likely pathway for visual interhemispheric transfer.

The source and target cortical areas of the visual interhemispheric transfer remain to be fully understood. Previous studies have demonstrated the existence of a small patch of transcallosal axons between visual areas 17 and 18 (Clarke and Miklossy, 1990; Aboitiz and Montiel, 2003). This motivated our choice of source and target cortical areas, which led to conduction velocity estimates consistent with previous studies (Caminiti et al., 2013). However, it has been suggested that these connections alone might be insufficient to produce an effective interhemispheric transfer (Innocenti et al., 2015) and higher visual processing areas might be involved.

### 5.4 Future prospects

The proposed model requires a measure of the MRI g-ratio and axonal conduction velocity in a specific white matter tract of interest. Since measurements of conduction velocity may only be conducted for a limited set of white matter tracts in the human brain, this model, in its current form, cannot be extended to the entire white matter, in contrast to other models (*e*.*g*. Assaf et al., 2008; Zhang et al., 2011). Instead, this model is geared towards the detailed characterization of a restricted set of white matter tracts. In its proposed form, two parameters relating to the morphology of white matter axons may be estimated from a total of four. Extending the number of estimated parameters requires the use of additional data integrated into the proposed framework. The nature of this data needs to be carefully considered. Alternatively, as was outlined in this work, a subset of the model parameters may be set to a constant value based on histological studies. A prime candidate is the mode of the axonal radius distribution (*M*), preserved between white matter tracts, individuals, and animal species (Tomasi et al., 2012; Liewald et al., 2014). This choice may need to be reconsidered for the study of brain pathologies that might differentially affect axons of different sizes.

In the current implementation, the estimate of the IHTT is calculated from the average of the EEG CD across participants. This leads to an average bias in *θ* and *β* is of 12% and 0.9% across participants, reaching up to 40% and 3% respectively (Fig. 6). While small compared to differences between tracts (∼350% for *θ* and ∼7% for *β*, see Fig. 4), this is of the order of the inter-subject differences for these parameters (Fig. 9) and prevents the estimation of axonal morphological features at the individual level of each dataset. This limitation represents the primary avenue for future implementations of our model. Estimation of the IHTT from alternative techniques such as transcranial magnetic stimulation should be considered (Lo and Fook-Chong, 2004; Spitzer et al., 2004; Basso et al., 2006; Deftereos et al., 2008; Marzi et al., 2009). Following these improvements, the question of the accuracy of the parameter estimates and their comparison with histological data might arise. We highlight that the parameters of the proposed model touch on properties of brain tissue that have received little attention in histological studies to date. These include the radius dependence of the axonal g-ratio and the radius dependence of fiber myelination change in health and disease. Validation of the proposed model may therefore bring opportunities for new research avenues for histological studies of the human brain.

## 6 Conclusions

In summary, we present a novel method that allows the estimation of morphological properties of axons from MRI and EEG data acquired in-vivo in healthy volunteers. This method enables the combined estimation of axonal radius and myelin thickness and opens the way for improved specificity in the studies of the brain conducted from in-vivo data. The method enables the assessment of the distribution of morphological features across axons and represents a significant step towards in-vivo histological studies in the human brain.

## 8 Author Contributions

RO conception and design of the study, acquisition of MRI and EEG data, data analysis and interpretation, drafting of the manuscript, and critical revision. AP acquisition of EEG data, data interpretation, and manuscript revision. GD acquisition of MRI data, data interpretation, and manuscript revision. MDL conception and design of the study, data interpretation, drafting of the manuscript, and critical revision. AL conception and design of the study, data interpretation, drafting of the manuscript, and critical revision.

## 10 Acknowledgments

The MRI data was acquired on the MRI platform of the Clinical Neuroscience Department, Lausanne University Hospital.

## 12 Data Availability Statement

This material will be made available to the general public upon publication of this manuscript.

