## Supplementary Material for "In-vivo estimation of axonal morphology from MRI and EEG data"

### 1 Supplementary Figures

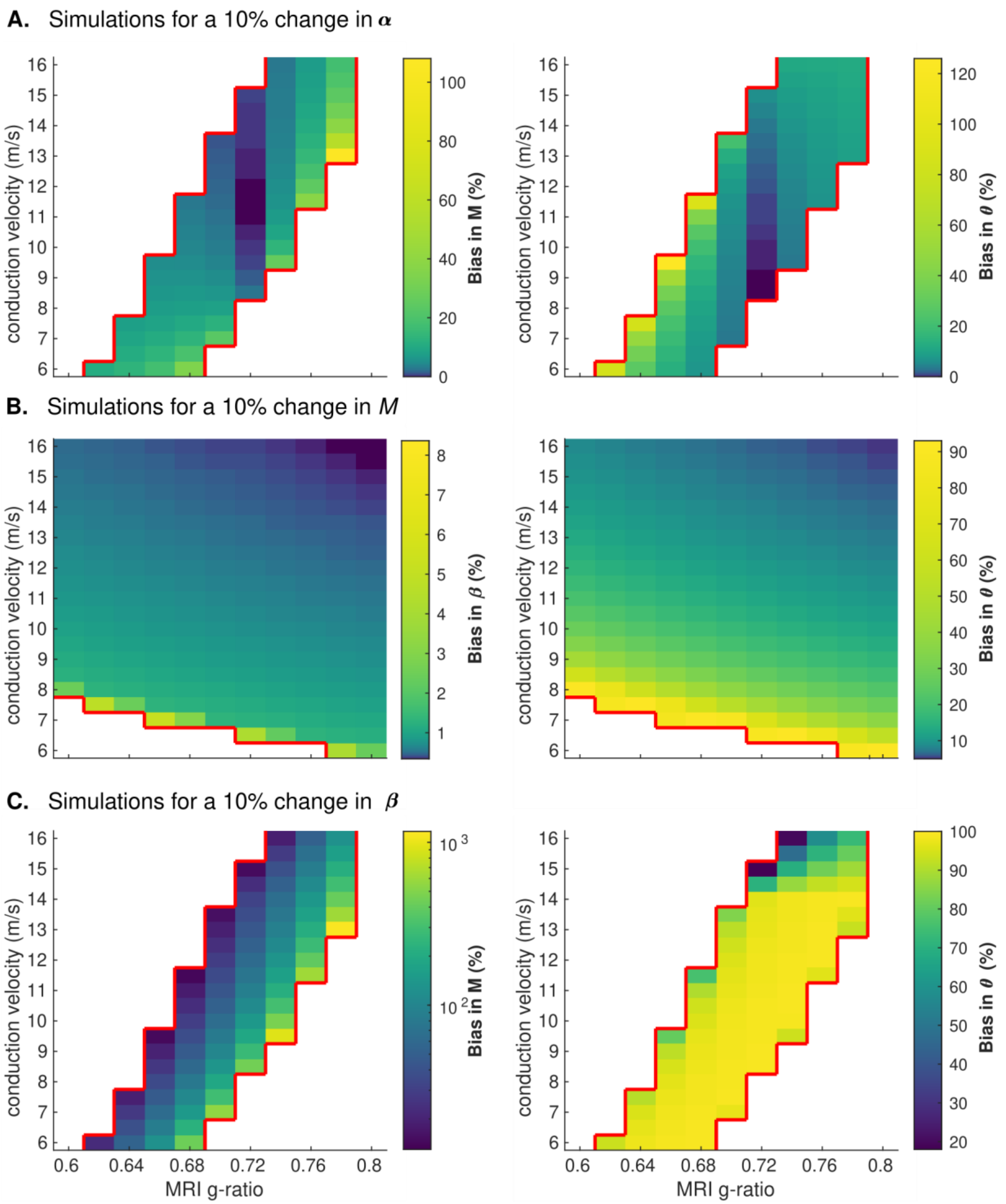


**Supplementary Figure 1.** Impact of inaccurate constant parameters on the estimated morphological features. (A) A 10% bias in α leads to an average bias of ~10% and ~15% for $M$ and $\theta$, respectively. For a representative combination of in-vivo parameter estimates ($g_{MRI}$=0.72, $V$=10 m/s), this bias is

~0.5% for $M$ and $\theta$. (B) A 10% bias of $M$ leads to an average bias of ~1% and ~22% for $\beta$ and $\theta$, respectively. For the representative combination of $g_{MRI}$ and $V$, this bias is <1% and ~16%. (C) A 10% bias of $\beta$ leads to an average bias of ~155% and ~92% for $M$ and $\theta$, respectively. For the representative combination of $g_{MRI}$ and $V$, this bias is ~145% for $M$ and ~100% for $\theta$. The red contour lines delineate the region of biologically plausible parameter values, compatible with the proposed model (Fig 3). Estimates outside this region were never observed in the in-vivo data and produced large values in these analyses and are therefore masked out.

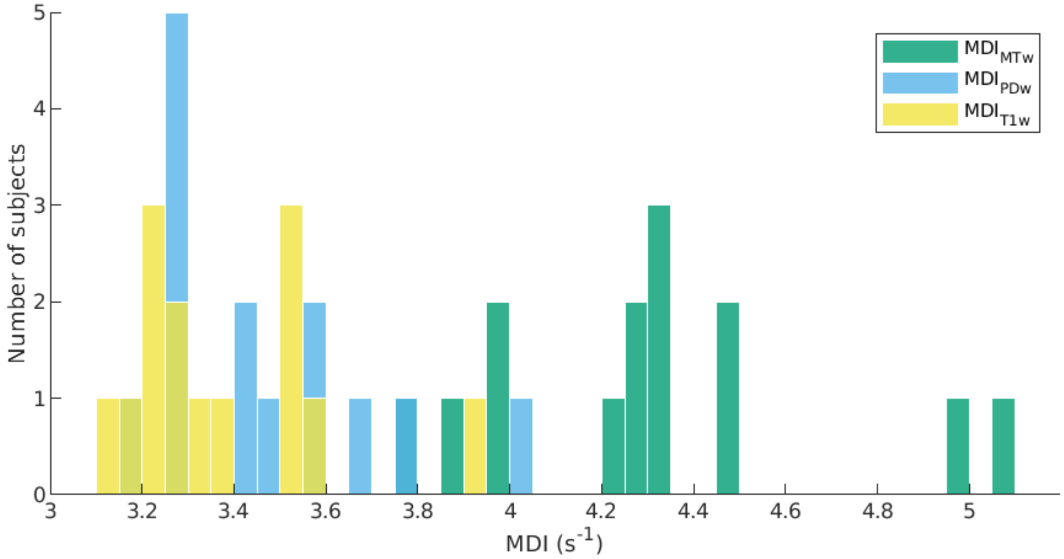


**Supplementary Figure 2.** Distribution of the Motion Degradation Index (MDI, $s^{-1}$) values for the raw FLASH images across all subjects (N=14).

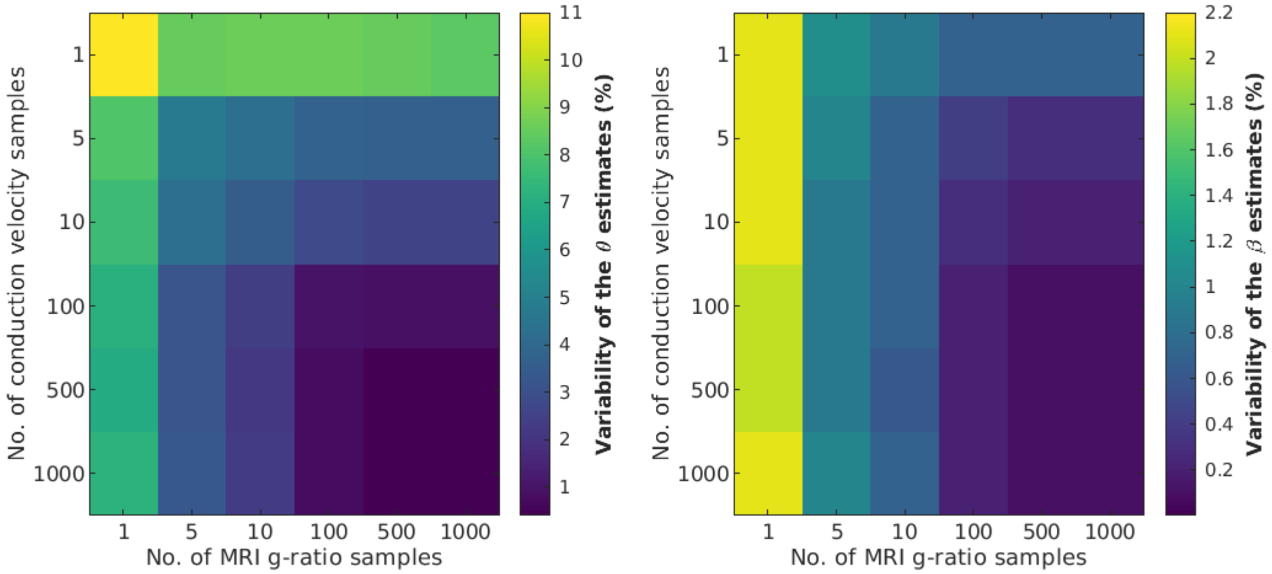


**Supplementary Figure 3.** Dependence of the variability of the $\theta$ (left) and $\beta$ (right) estimates on the number $g_{MRI}$ and $V$ samples. The variability of the $\theta$ estimates shows a strong dependence on the number of samples of both $V$ and $g_{MRI}$. The variability of $\beta$ is mostly driven by the number of $g_{MRI}$ samples.

### 2 Supplementary Appendix A

Our two morphological features of interest are defined as (see Theory section):

$$g(r) = \beta * r^{\alpha}$$

$$P(r) = P(r|k,\theta) = \frac{1}{\Gamma(k)\theta^k} r^{k-1} e^{\frac{-r}{\theta}}$$

With $k$ being the shape and $\theta$ the scale of the axonal radius distribution.
In agreement with West et al., 2016, the MRI g-ratio is written as an ensemble average of the axonal g-ratios within each image voxel, weighted by the axons' cross-sectional area:

$$g_{MRI}{}^2 = \frac{\int_0^{\infty} R^2 g(r)^2 P(r)\,dr}{\int_0^{\infty} R^2 P(r)\,dr} = \frac{\int_0^{\infty} r^2 P(r)\,dr}{\int_0^{\infty} \frac{r^2}{g(r)^2} P(r)\,dr} \tag{1}$$

where $r$ is the axon radius and $R$ the fiber radius ($R = r/g(r)$).

Using $P(r)$ and $g(r)$ as we defined previously, the integrand in the denominator can be re-written as a Gamma distribution with shape parameter $k' = k - 2\alpha$, leading to:

$$g_{MRI}{}^2 = \frac{\beta^2 \int_0^{\infty} r^2 P(r)\,dr}{\int_0^{\infty} r^{2-2\alpha} P(r)\,dr} = \frac{\beta^2 \int_0^{\infty} r^2(r) P(r|k,\theta)\,dr}{\frac{\Gamma(k')\theta^{k'}}{\Gamma(k)\theta^k} \int_0^{\infty} r^2 P(r|k',\theta)\,dr} \tag{2}$$

These integrals can be transformed using the second moment of the radius distribution:

$$g_{MRI}{}^2 = \frac{\beta^2 \Gamma(k)\theta^k}{\Gamma(k')\theta^{k'}} \frac{(k+1)k\theta^2}{(k'+1)k'\theta^2} = \frac{\beta^2 \Gamma(k)\theta^{2\alpha}}{\Gamma(k')} \frac{(k+1)k}{(k'+1)k'} \tag{3}$$

For convenience, we make a change of variable, where $M = \theta(k-1)$, corresponding to the mode/peak of the radius distribution. Eq. (3) can be re-written as:

$$g_{MRI}{}^2 = \beta^2 \theta^{2\alpha} * \frac{\Gamma\left(\frac{M}{\theta} + 1\right)}{\Gamma\left(\frac{M}{\theta} + 1 - 2\alpha\right)} \frac{M + 3\theta + 2\theta^2/M}{M + (3 - 4\alpha)\theta + (2 + 4\alpha^2 - 6\alpha)\,\theta^2/M} \tag{4}$$

As described in Waxman and Bennett, 1972, axonal conduction velocity ($v$) can be derived from the morphological properties of axons using: $v\,[m/s] = p * \frac{d\,[\mu m]}{g}$, where $p$ (~5.5-6.0) represents the contribution of additional axonal factors to the propagation of action potentials (*e.g.* length of Ranvier nodes, electrical properties of the myelin membranes). Assuming an equal contribution from all axons to the conduction velocity $V$ measured with EEG, we obtain:

$$V = 5.5 \int_0^{\infty} \frac{2r\,P(r)}{g(r)}\,dr \tag{5}$$

Using $P(r)$ and $g(r)$ as we defined previously, the integrand in the denominator can be re-written as a Gamma distribution with shape parameter $k" = k - \alpha$, leading to:

$$V = \frac{11}{\beta} \int_0^{\infty} r^{1-\alpha} P(r|k,\theta)\,dr = \frac{11}{\beta} \frac{\Gamma(k'')\theta^{k''}}{\Gamma(k)\,\theta^k} \int_0^{\infty} rP(r|k'',\theta)\,dr \tag{6}$$

These integrals can be transformed using the first moment of the radius distribution:

$$V = \frac{11}{\beta} \frac{\Gamma(k'')\theta^{k''}}{\Gamma(k)\theta^{k}} k''\theta \tag{7}$$

This expression can be re-written as a function of $M$ and $\theta$:

$$V = \frac{11 * \theta^{1-\alpha}}{\beta} \frac{\Gamma\left(\frac{M}{\theta} + 1 - \alpha\right)}{\Gamma\left(\frac{M}{\theta} + 1\right)} \left(\frac{M}{\theta} + 1 - \alpha\right) \tag{8}$$

Our final model incorporates Eqs. (4) and (8).
